# Identification of biomarkers and candidate inhibitors for multiple myeloma

**DOI:** 10.1101/2021.02.25.432847

**Authors:** Hanming Gu, Wei Wang, Gongsheng Yuan

**Author notes:** Corresponding author: Dr. Gongsheng Yuan, Department of Physiology and Pathophysiology, School of Basic Medical Sciences, Fudan University, Shanghai, China.

## Abstract

Multiple myeloma (MM) is a plasma cell malignancy that causes the overabundance of monoclonal paraprotein (M protein) and organ damages. In our study, we aim to identify biological markers and processes of MM using a bioinformatics method to elucidate their potential pathogenesis. The gene expression profiles of the GSE153626 datasets were originally produced by using the high-throughput Illumina HiSeq 4000 (Mus musculus). The functional categories and biochemical pathways were identified and analyzed by the Kyoto Encyclopedia of Genes and Genomes pathway (KEGG), Gene Ontology (GO), and Reactom enrichment. KEGG and GO results showed the biological pathways related to immune dysfunction and signal transduction are mostly affected in the development of MM. Moreover, we identified several genes including Gngt2, Foxp3, and Cd3g were involved in the regulation of immune cells. We further predicted new inhibitors that have the ability to block the progression of MM based on the L1000fwd analysis. Therefore, this study provides further insights into the underlying pathogenesis of MM.

## Introduction

Multiple myeloma is hematological cancer that caused end-organ damage including anemia, renal impairment, and bone lesions^1^. MM primarily occurs in the elderly (over 65 years) and the global incidence has increased dramatically since 1990^2, 3^. With current chemotherapy and regimens, the survival rate has improved, although it is still a high-risk disease^4^. Malignant migration of MM is a consequence of a combination of factors such as genetic mutations and genetic evolutions in the BM environment^5^. Moreover, immune dysfunction was found in myeloma patients, raising the question of whether immunological dysregulation is an important mechanism of disease development^6, 7^.

Most of the findings showed that several types of mouse models have been used, which aim to provide insights on the relationship between MM and BM microenvironment^8^. Here, the mouse model we analyzed was created through conditionally activating the endogenous NrasQ61R and an MYC transgene in germinal center B cells^9^. This mouse model developed a classic malignant MM characterized by phenotype, signaling pathways, and MM gene signatures^9^. Recent studies suggest that the uncontrolled immunological microenvironment leads to the progress of MM^10^. The activated MM may cause bone damage, anemia, multiple organ metastasis^11^. However, the mechanism underlying MM is still unknown.

Here, we studied the relative changes of genes in MM mouse models. Then, we identified and analyzed a series of DEGs, the relevant biological process, and potential drug targets of MM using comprehensive bioinformatic analysis. Last, we performed the gene and pathway analysis, the functional enrichment, and protein-protein interaction (PPI) for discovering MM gene nodes. The critical genes and pathways could be in favor of future clinical and therapeutic studies.

## Methods

### Data resources

The dataset GSE153626 was downloaded from the GEO database (http://www.ncbi.nlm.nih.gov/geo/). The data was produced by Illumina HiSeq 4000 (Mus musculus), Blood Research Institute, Versiti 8727 W Watertown Plank Rd, Milwaukee, Wisconsin. Bulk RNA-Seq analysis was performed using CD138+ B220− CD45.2+ bone marrow cells of three control and five MM models conditionally activating expression of endogenous NrasQ61R and a MYC transgene in germinal center B cells (VQ mice).

### Data acquisition and preprocessing

The dataset GSE153626 containing WT samples and VQ mice samples was analyzed and conducted by R script as described previously^12, 13^. We used a classical t test to identify DEGs with P<.01 and fold change ≥1.5 as being statistically significant.

### Gene functional analysis

Gene ontology (GO) analysis is a widely used approach to develop a comprehensive and computational model of biological systems to analyze genomic data and define characteristic biological information^14^. The Kyoto Encyclopedia of Genes and Genomes (KEGG) database is commonly used for understanding high-level functions and utilities of the biological system^15^. We performed the GO analysis and KEGG pathway enrichment analysis by using the Database for Annotation, Visualization, and Integrated Discovery (DAVID) (http://david.ncifcrf.gov/) as described previously^16, 17^. P<.05 and gene counts >10 were considered statistically significant.

### Module analysis

The Molecular Complex Detection (MCODE) of cytoscape software was used to analyze the densely connected regions in protein-protein interaction (PPI) networks^18^. The significant modules were from the constructed PPI network using MCODE^19^. The function and pathway enrichment analyses were performed by using DAVID, and P<.05 was used as the cutoff criterion.

### Reactome pathway analysis

The dataset was performed by the Reactom pathway (https://reactome.org/) to obtain the visualization, interpretation and analysis of potential pathways. P<.05 was considered statistically significant.

## Results

### Identification of DEGs in myeloma cells from MM models

Multiple myeloma mouse models were created by activating the expression of endogenous NrasQ61R and a MYC transgene in germinal center B cells (VQ mice) at Blood Research Institute, Versiti (Wisconsin, USA). To gain the insights on MM model genes, the modular transcriptional signature of VQ mice was compared to that of WT mice. A total of 3721 genes were identified to be differentially expressed in MM models with the threshold of P<0.005. The top 10 up- and down-regulated genes for VQ mice and WT control samples are listed in Table 1.

**Table 1.**
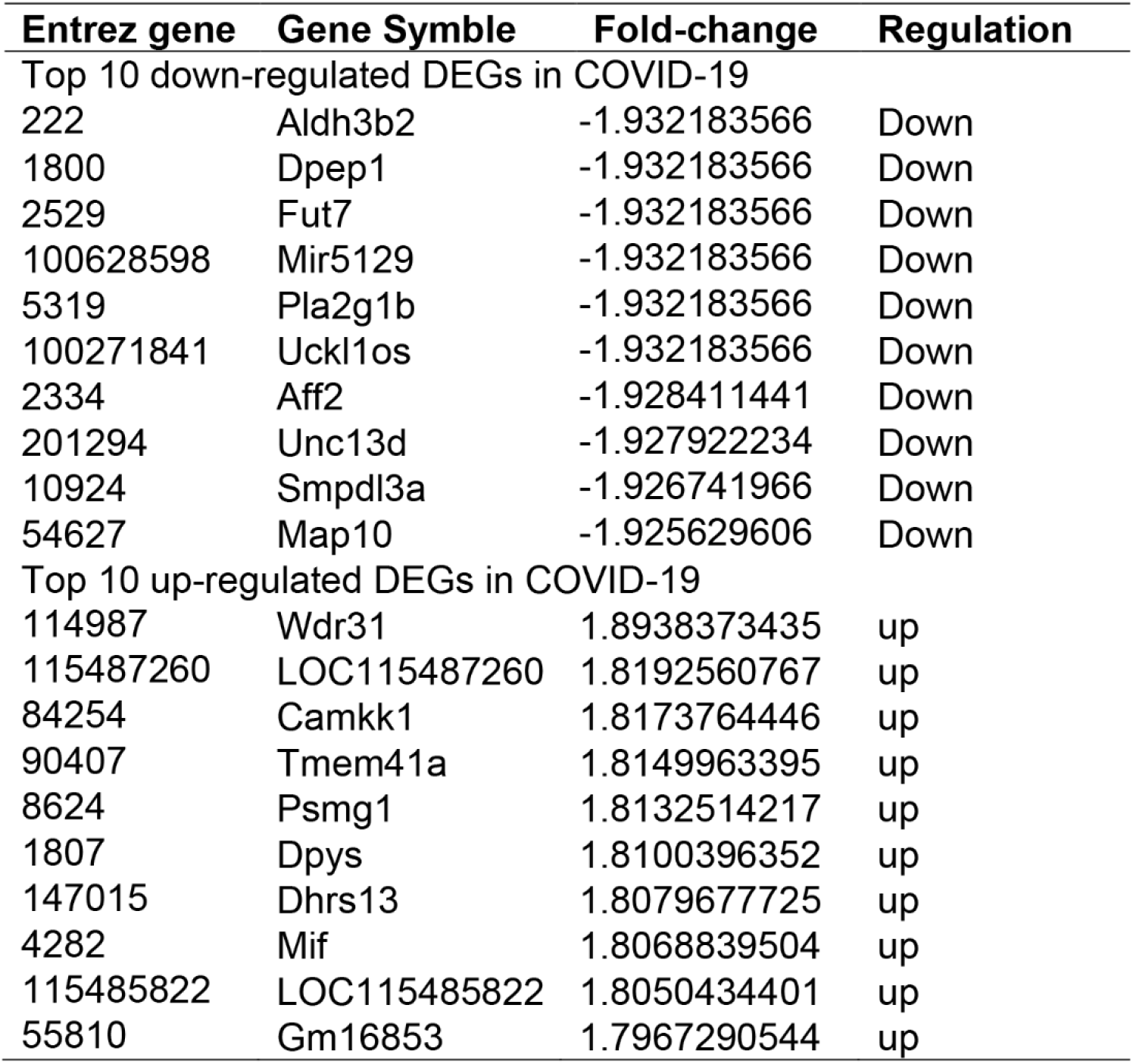

### KEGG analysis of DEGs in MM models

To further identify the biological roles and potential mechanisms of the DEGs from the MM models versus WT controls, we performed KEGG pathway and GO categories enrichment analysis (Supplemental Table S1). KEGG pathways (http://www.genome.jp/kegg/) are a collection of manually drawn pathway maps for understanding the molecular interaction, reaction and relation networks. Our study presented the top ten enriched KEGG pathways including “Inflammatory bowel disease (IBD)”, “Hematopoietic cell lineage”, “African trypanosomiasis”, “Transcriptional misregulation in cancer”, “Pancreatic secretion”, “Fc epsilon RI signaling pathway”, “T cell receptor signaling pathway”, “Natural killer cell mediated cytotoxicity”, “Salivary secretion”, and “Oxytocin signaling pathway” (Figure 1).

**Figure 1.**
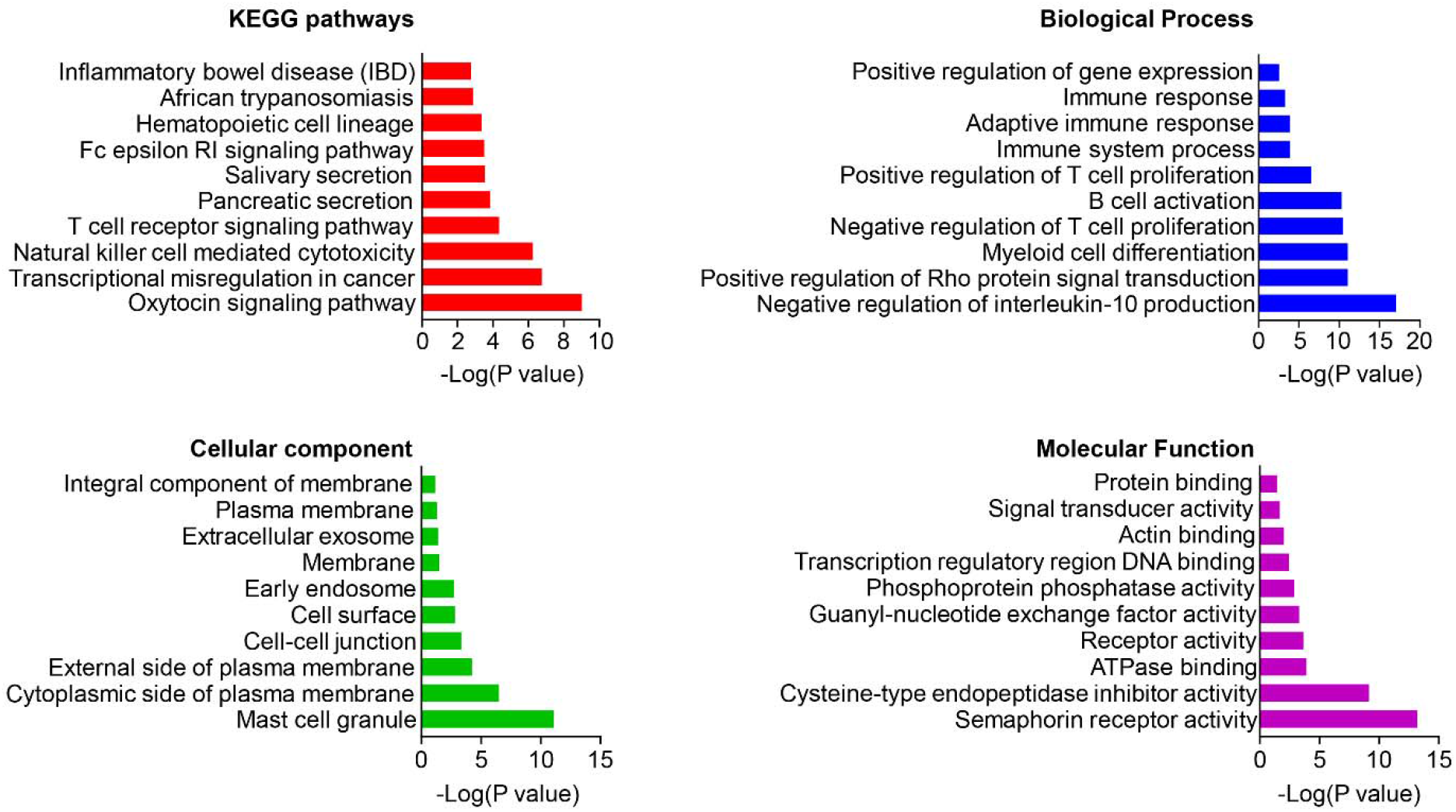
The KEGG pathways, biological process, cellular component, and molecular function terms enriched by the DEGs between WT and MM mouse models. DEGs =differentially expressed genes, KEGG = Kyoto Encyclopedia of Genes and Genomes.

### GO analysis of DEGs in MM models

Gene ontology (GO) analysis is a system for hierarchically classifying genes, which includes cellular components (CC), molecular functions (MF), and biological processes (BP)^20^. Here, we identified top ten cellular components including “Membrane”, “External side of plasma membrane”, “Cell surface”, “Plasma membrane”, “Cell-cell junction”, “Mast cell granule”, “Extracellular exosome”, “Cytoplasmic side of plasma membrane”, “Early endosome”, and “Integral component of membrane” (Figure 1). We then identified top ten biological processes: “Negative regulation of interleukin-10 production”, “Positive regulation of Rho protein signal transduction”, “Myeloid cell differentiation”, “Negative regulation of T cell proliferation”, “B cell activation”, “Positive regulation of T cell proliferation”, “Immune system process”, “Adaptive immune response”, “Immune response”, and “Positive regulation of gene expression” (Figure 1). Last, we identified top ten molecular functions: “Protein binding”, “Receptor activity”, “Guanyl-nucleotide exchange factor activity”, “Cysteine-type endopeptidase inhibitor activity involved in apoptotic process”, “ATPase binding”, “Semaphorin receptor activity”, “Transcription regulatory region DNA binding”, “Phosphoprotein phosphatase activity”, “Actin binding”, and “Signal transducer activity” (Figure 1).

### PPI network and Module analysis

The PPI networks were created to analyze the relationships of DGEs at the protein level^21, 22^. The criterion of combined score >0.7 was chosen and the PPI network was constructed by using the 143 nodes and 209 interactions. Among these nodes, the top ten genes with highest scores are shown in Table 2. The top two significant modules of MM models versus WT samples were selected to indicate the functional annotation (Figure 2).

**Table 2.** Top ten genes demonstrated by connectivity degree in the PPI network.

| Gene Symbol | Gene title | Degree |
| --- | --- | --- |
| Gngt2 | G protein subunit gamma transducin 2 | 15 |
| Foxp3 | Forkhead box P3 | 12 |
| Cd3g | CD3g molecule | 10 |
| H2-Ab1 | Histocompatibility 2, class II antigen Ab1 | 9 |
| H2-Eb2 | Histocompatibility 2, class II antigen Eb2 | 8 |
| F2rl2 | Coagulation factor II thrombin receptor like 2 | 8 |
| Ccr5 | C-C motif chemokine receptor 5 | 7 |
| Ceacam1 | CEA cell adhesion molecule 1 | 7 |
| Vav3 | Vav guanine nucleotide exchange factor 3 | 7 |
| Lat | Linker for activation of T cells | 7 |

**Figure 2.**
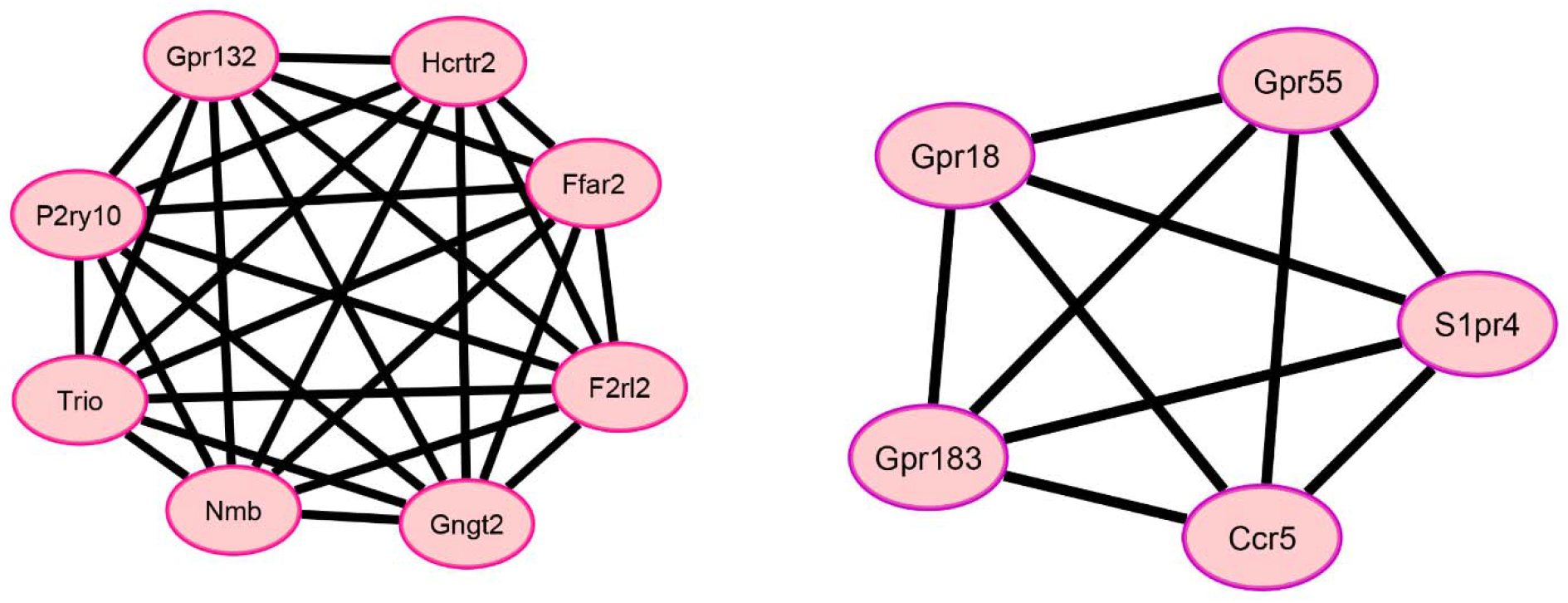
Top two modules from the PPI network between WT and MM mouse models.

### Reactome Pathway in MM models

We identified several signaling pathways by using Reactome Pathway Database (https://reactome.org/). We identified top ten signaling pathways including: “Interleukin-35 Signalling”, “RUNX1 and FOXP3 control the development of regulatory T lymphocytes (Tregs)”, “Other semaphorin interactions”, “TFAP2 (AP-2) family regulates transcription of growth factors and their receptors”, “Interleukin-18 signaling”, “Neurophilin interactions with VEGF and VEGFR”, “Defective factor IX causes hemophilia B”, “Interleukin-12 family signaling”, “Interleukin-38 signaling”, and “RUNX1 regulates transcription of genes involved in BCR signaling” (Supplemental Table S2). We then constructed the reaction map according to the signaling pathways (Figure 3).

**Figure 3.**
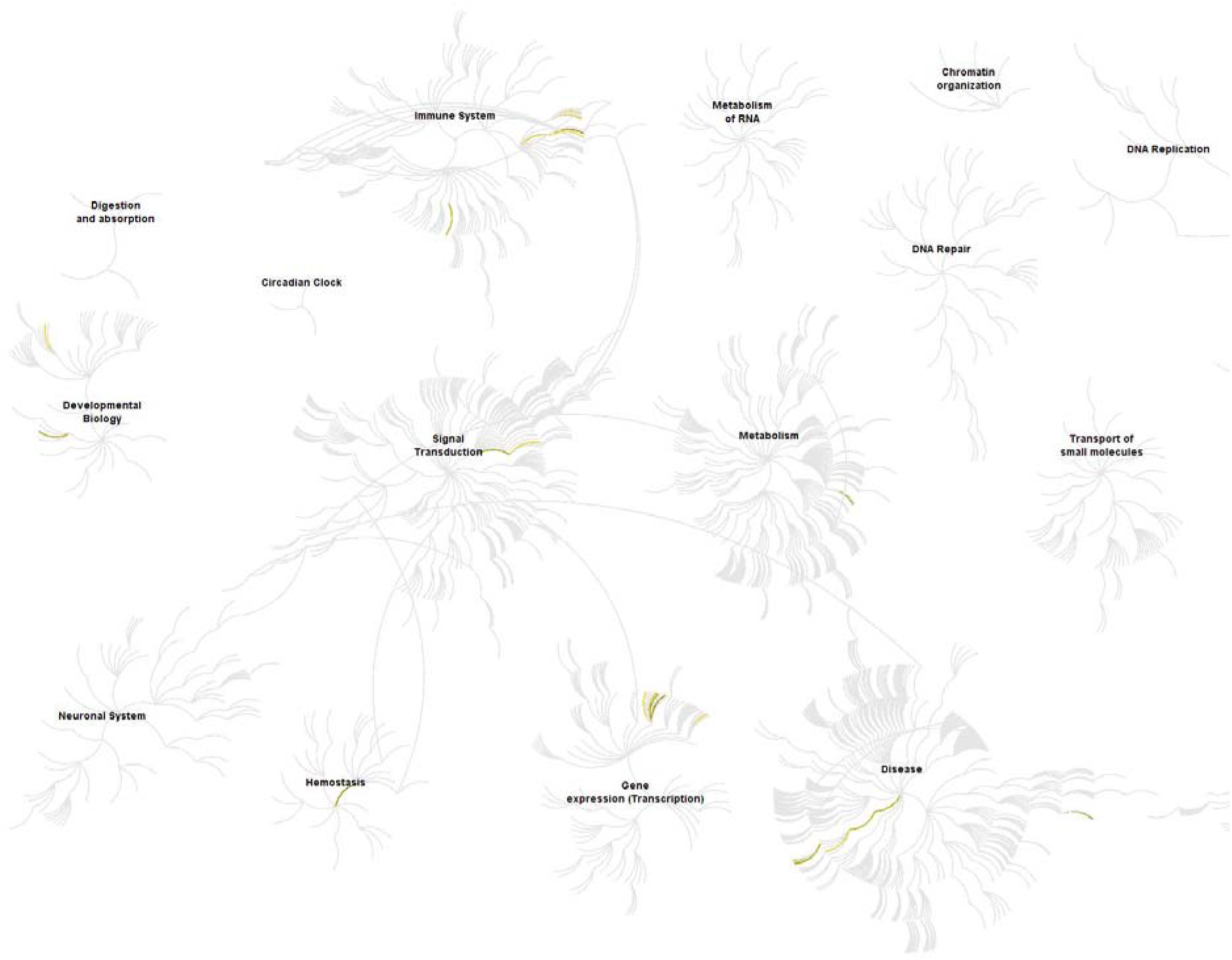
The Reactom pathway visualization. Input genes are represented by the top significantly changed genes obtained from the GSE153626 dataset (P <0.01). The yellow color represents the most relevant signaling pathways.

### Potential inhibitors in MM models

To further know how to prevent the MM, we introduced the L1000FDW system that can predict and analyze the potential inhibitors (Figure 4). With the mechanisms-of-action (MOA), the L1000FDW system indicated the most potentially affected pathways. We selected top ten inhibitors according to the DEGs and the inhibitor maps: “BRD-A47816767”, “radicicol”, “melperone”, “IKK-2-inhibitor-V”, “I-606051”, “alitretinoin”, “BRD-K70771662”, “SSR-69071”, “SA-1447005”, and “CAY-10470” (Supplemental Table S3).

**Figure 4.**
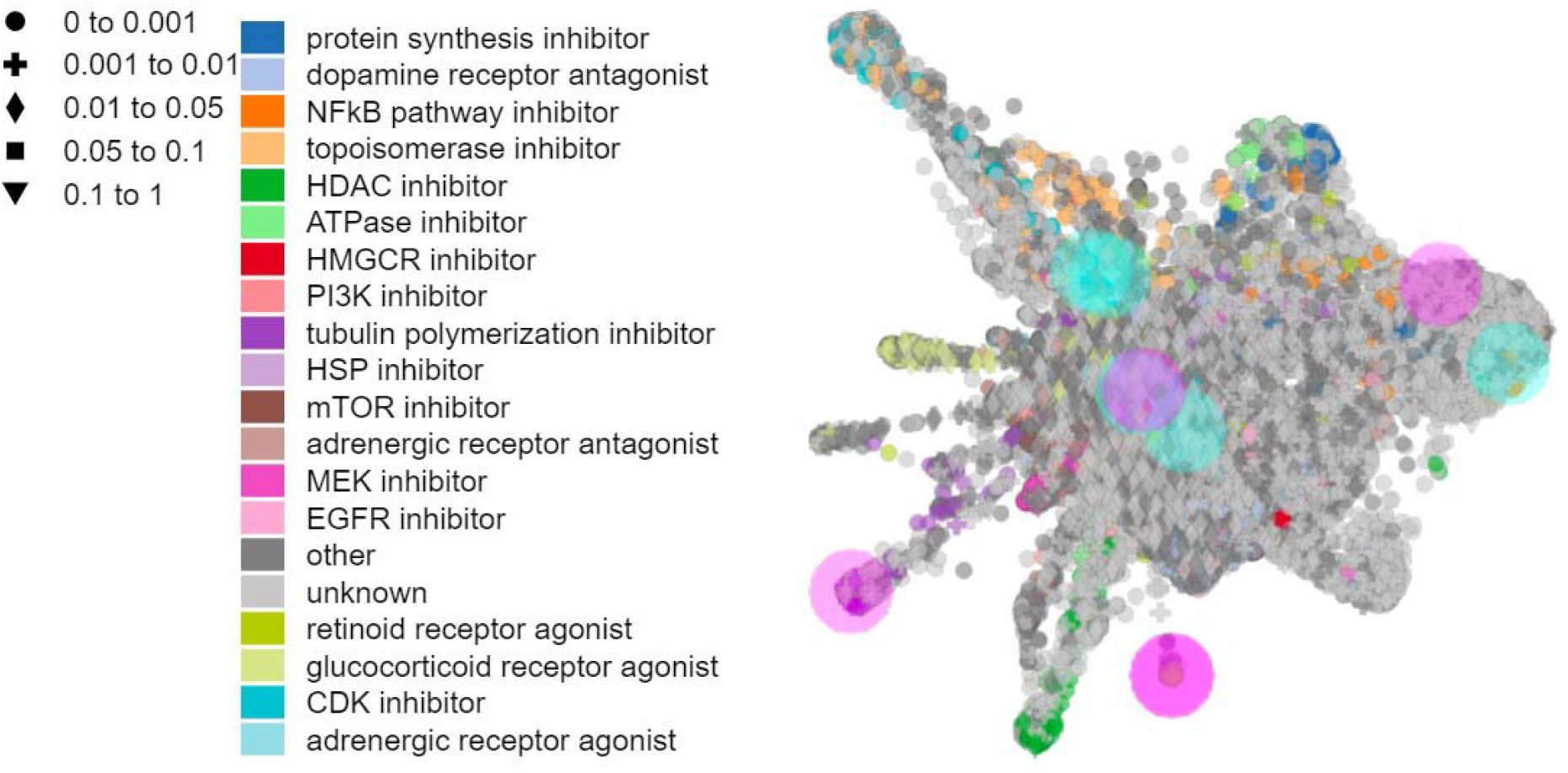
Inhibitor prediction against MM by L1000FDW visualization. Input genes are represented by the significantly changed genes obtained from the GSE153626 dataset. Dots are the Mode of Action (MOA) of the respective drug.

## Discussion

MM is a plasma cell malignancy that accounts for about 10% of hematologic cancers^23^. MM causes several symptoms such as organ damage, anemia and lytic bony lesions^24^. MM was triggered from a premalignant state called “monoclonal gammopathy of undetermined significance (MGUS)” ^25^. Malignant transformation is the result of combinations of multiple factors such as genes and BM microenvironment^26^. Moreover, immune dysfunction plays an important role in the progression of MM^6^. Thus, the signature of genes and proteins are the key targets to prevent the diseases.

To better understand the effects of inhibitors on the MM mouse models, we analyzed a novel MM mouse model that was created by Blood Research Institute, Versiti. This model was based on the conditionally activating expression of endogenous NrasQ61R and a MYC transgene in germinal center B cells (VQ mice)^9^. Thus, by using this transgenic mouse we could produce more MM models to test the drugs and inhibitors as previously described^17^. By analyzing the DEGs of this model, we selected 10 proteins that may be important during the development of MM according to the PPI network analysis. G proteins and RGS proteins are widely involved in the immune and inflammatory diseases. In our study, the Guanine nucleotide-binding protein G(I)/G(S)/G(O) subunit gamma-T2 is a potential prognostic marker of esophageal cancer^27^. The FOXP3 in T-cells can suppress antigen priming of lymphocytes and can be regulated by the multifunctional circadian clocks^28–30^. CD3g deletion from a consanguineous family is associated with autoimmunity^31^. H2-Ab1 and H2-Eb2 are reported to be involved in the control of tuberculosis infection^32^. G-protein-coupled receptor (GPCR) proteins and RGS proteins are highly related to the tumor genesis^33–36^. F2Rl2 is recently reported to be a GPCR that encodes protease-activated receptor-3 (PAR3)^37^. Circadian clock controls numerous genes to regulate the cell functions during normal conditions and diseases^29, 38–41^. As a clock-controlled gene, CCR5 is reported to suppress anti-tumor immune response and promote tumor growth^42, 43^. Long and short isoforms of CEACAM1 are strongly expressed on cancer cells in the liver metastases^44^. VAV3 is associated with the ERBB4-mediated cancer cell migration^45^. LAT-1 is identified as a promotor in gastric cancer that is related to the clinicopathologic features^46^.

KEGG and GO analysis showed that immune responses play a critical role in the progression of MM. The KEGG analysis showed the “Natural killer cell mediated cytotoxicity”, “T cell receptor signaling pathway”, “Fc epsilon RI signaling pathway” and “Inflammatory bowel disease (IBD)” were the major pathways during the development of MM. It is suggested that the occurrence of MM is based on the inflammation or immune changes. Similarly, MM is reported to be caused by the disrupted immune surveillance such as deregulation of the T and natural killer cell compartment and recruitment of immunosuppressive cells^47^. Interestingly, the BP of GO analysis showed “Negative regulation of IL10 production”, “Negative regulation of T cell proliferation” and “B cell activation”, suggesting that the multiple immune cells were involved in the progression of MM not only B cells. Thus, the BM microenvironmental condition is crucial in MM. Indeed, the MM cells disseminate in the BM microenvironment via interactions of the adhesion molecules and extracellular matrix (ECM) components and receive various signals that maintain their survival and activity^48^.

Despite recent advancements in drug development, the anti-cancer drug resistance is a major limitation of MM therapy. Thus, the potential inhibitors may be a good substitute for the traditional drugs of MM. In our study, we predicted several inhibitors of MM that most of them are involved in the regulation of immune and protein synthesis pathways such as NF-κB pathways. NF-κB pathway is a center of inflammation which is widely associated in the regulations and controls of diseases^36, 49^. However, the side effect of NF-κB inhibitors is so strong that they cannot be used in clinical^50^. Here, besides NF-κB inhibitors, we selected several novel inhibitors that may take effects on the MM mouse models. Our future studies will mainly target the effects on the treatment of MM by using MM mouse models or primary MM cells.

In summary, we identified the potential biomarkers for MM. Immune dysfunction and BM microenvironment are two key processes in the novel MM mouse models. Future studies will focus on the administration of potential inhibitors on clinical trials. This study thus provides further insights into the mechanism of MM, which may facilitate the diagnosis and drug development.

